# Lipid composition but not curvature is a determinant of a low molecular mobility within HIV-1 lipid envelope

**DOI:** 10.1101/315168

**Authors:** Iztok Urbančič, Juliane Brun, Dilip Shrestha, Dominic Waithe, Christian Eggeling, Jakub Chojnacki

## Abstract

Human Immunodeficiency Virus type-1 (HIV-1) acquires its lipid membrane from the plasma membrane of the infected cell from where it buds out. Previous studies have shown that the HIV-1 envelope is a very low mobility environment with the diffusion of incorporated proteins two orders of magnitude slower than in plasma membrane. One of the reasons for this difference is thought to be due to HIV-1 membrane composition that is characterised by a high degree of rigidity and lipid packing. To further refine the model of the molecular mobility on HIV-1 surface, we here investigated the relative importance of membrane composition and curvature in Large Unilamellar Vesicles of different composition and size (0.2–1 μm) by super-resolution STED microscopy-based fluorescence correlation spectroscopy (STED-FCS) analysis. We find that lipid composition and its rigidity but not membrane curvature play an important role in the decreased molecular mobility on vesicle surface thus confirming that this factor is an essential determinant of HIV-1 low surface mobility. Our results provide further insight into the dynamic properties of the surface of individual HIV-1 particles.

## 1. Introduction

Human Immunodeficiency Virus Type-1 (HIV-1) is an enveloped retrovirus. It acquires its lipid membrane from the plasma membrane of the infected cell during the budding process driven by the assembly of the viral structural protein Gag [1]. In the budded, morphologically mature HIV-1 particle (Figure 1 a) this combination of lipids, viral structural proteins as well as membrane incorporated viral fusion protein Env and other cellular proteins create a unique lipid/protein surface environment that is highly curved due to the size ( <140 nm) of the virus particle. Lipidomic studies of isolated viral lipids have shown that in comparison to the plasma membrane HIV-1 membrane is enriched in sphingomyelins (SMs), glycosphingolipids, cholesterol (Chol) and phosphoinositides such as phosphatidylinositol (4,5) bisphosphate lipid (PIP2) [2,3]. Such an environment is characterised by a high degree of lipid packing and therefore low polarity within the lipid bilayer. When such membrane is studied with polarity-sensing probes such as Laurdan this results in a blue-shifted emission spectrum of the probe [4]. Such spectral changes are often compacted into the General Polarization (GP) parameter, spanning values between −1 and 1: high GP values indicate the existence of rigid, highly packed membranes, whereas low GP signifies fluid, less-packed lipid environments. Laurdan-based spectrophotometric studies of bulk purified particles [5] or spectral scanning microscopy-based analysis of individual particles [6] have indeed shown that HIV-1 membranes have a very high GP value of ~0.5, indicating a very high level of rigidity that has also been reported in highly cholesterol and sphingomyelin enriched model membranes [7] or giant plasma membrane vesicles [8]. All of this data supports the idea that HIV-1 may acquire its lipids and bud from the pre-existing so called “lipid raft” domains. On the other hand, a recent study on supported lipid bilayers suggested that instead of budding from existing lipid rafts Gag may instead create its own specialised lipid environment, by selectively trapping cholesterol and PIP2 at virus assembly sites [9].

**Figure 1.**
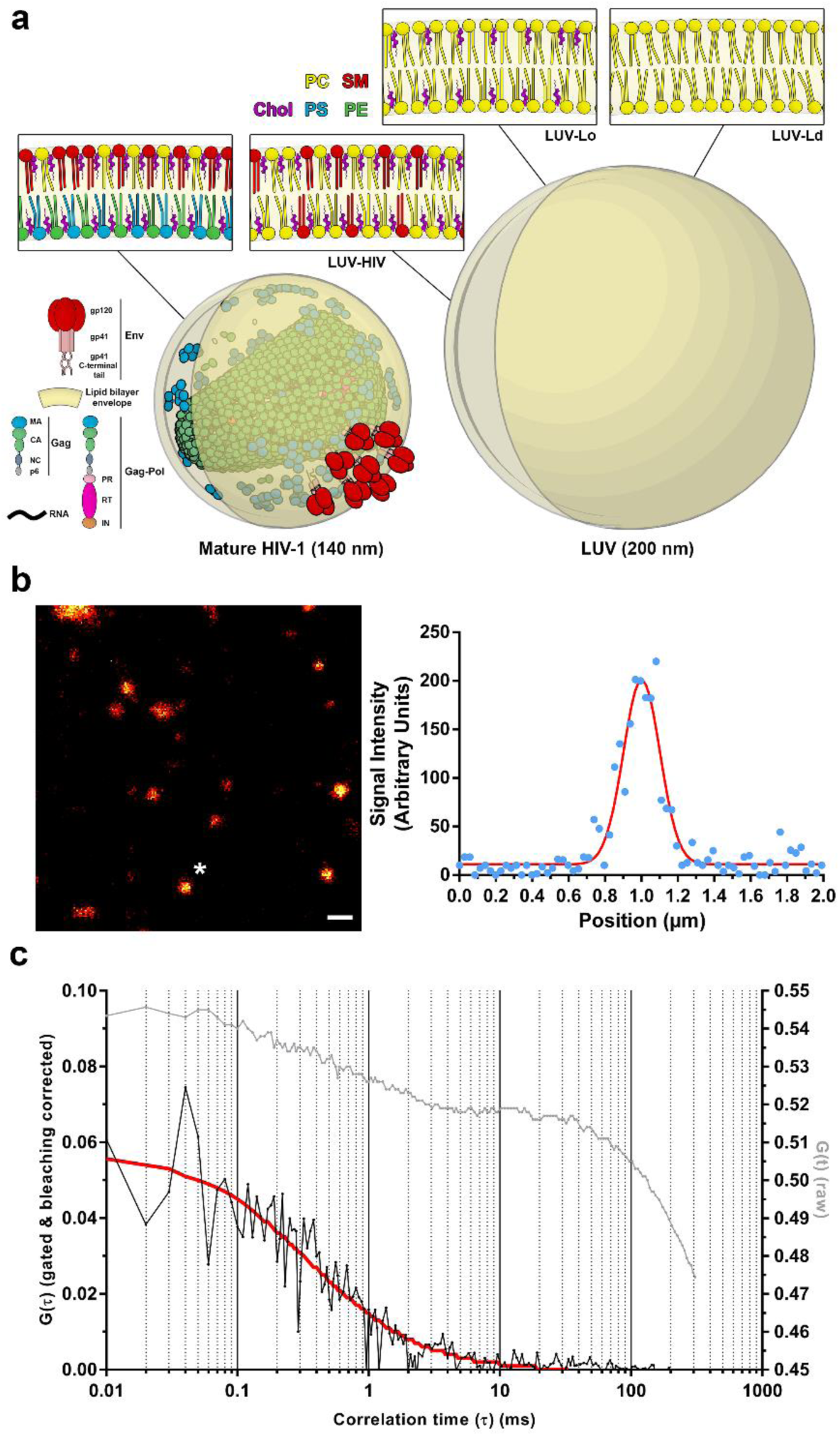
STED-FCS measurements of lipid diffusion in Large Unilamellar Vesicles (LUVs). (**a**) Schematic illustration of lipid and protein composition in mature HIV-1 (with Env surface and Gag matrix proteins and according sub-units) and 200 nm Large Unilamellar Vesicle (LUV) containing: DOPC (LUV-Ld), DOPC:Chol at 67:33 molar ratio (LUV-Lo) or DOPC:Chol:SM at 37:46:17 molar ratio (LUV-HIV). HIV-1 lipid composition was adopted from [3]. (**b**) Representative STED microscopy image of 200 nm LUV-Ld preparation. Imaging was used to locate LUVs of comparable size and brightness followed by the acquisition of the fluorescence fluctuation data. White star marks a 200 nm LUV used for analysis with its lateral fluorescence intensity profile shown on the right (blue points – raw intensity data, red line – fitted Gaussian curve). Scale bar: 500 nm; (**c**) Representative raw (grey) and gated & bleaching corrected (black) autocorrelation curves obtained from 200 nm LUV fluorescence fluctuation data. Gated & bleaching corrected autocorrelation curves were fitted using generic 2D diffusion model (red).

Despite this knowledge of overall characteristics of HIV-1 envelope, very little is known about the dynamic characteristics of this environment such as mobility of the individual molecules on the surface of individual particles. This is due to the fact that such measurements would require high spatial and temporal resolution to measure the diffusion of molecules confined within < 140 nm diameter of the virus particles, which is well below the diffraction limit of a conventional optical microscope. Thanks to recent advances in the field of super-resolution microscopy, techniques such as Stimulated Emission Depletion (STED) microscopy can now reach the desired sub 100 nm resolution that is required to image subviral structures [10]. Furthermore the combination of STED with high temporal resolution spectroscopic techniques such as Fluorescence Correlation Spectroscopy (FCS), allows determination of the molecular diffusion coefficients in the subdiffraction sized observation spots [11,12]. We have recently employed this technique to study the mobility of Env proteins on the surface of individual virus particles [6]. This study has established that the HIV-1 envelope is an intrinsically highly immobile environment where Env and other molecules such as major histocompatibility complex class-I (MHC-I) and glycophosphatidylinositol (GPI)-anchored proteins all exhibit a very low diffusion coefficient (*D* ≈ 0.002 μm^2^/s). This mobility is two orders of magnitude lower than mobility of these proteins when present in the cell plasma membrane [6]. Such a low mobility appears to be due to the combination of highly ordered and highly curved nature of the viral lipid envelope, tight packing of internal membrane-interacting virus proteins (MA domain of Gag) as well as passive incorporation of many types of cellular proteins during virus budding. Interestingly, the mobility is very low both in immature and mature viruses, despite the differences in Gag organisation (in the case of Env, mobility is even further decreased in immature viruses due to its tight links with the immature Gag shell) [6].

Here, we further refine the model of the molecular mobility on the virus surface by examining the factors that would play significant role in the reduction of molecular mobility. By measuring the lipid mobility on the surface of vesicles of different sizes and compositions characterised by different lipid packing we find that lipid composition and packing but not the membrane curvature plays an important role in the very low mobility of molecules on the HIV-1 surface.

## 2. Materials and Methods

### Lipids

Lipids 1,2-dioleoyl-sn-glycero-3-phosphocholine (DOPC), cholesterol (Chol), egg sphingomyelin (SM), and 1,2-distearoyl-sn-glycero-3-phosphoethanolamine-N-[biotinyl(polyethylene glycol)-2000] (DSPE-PEG-biotin) were purchased from Avanti Polar Lipids. The fluorescent lipid analogues Atto647N-DPPE and C-Laurdan were purchased from Atto-Tec and 2pprobes, respectively. Bovine serum albumin (BSA) and biotinylated BSA were obtained from Sigma Aldrich, whereas streptavidin was obtained from Life Technologies.

### Preparation of Large Unilamellar Vesicles (LUVs)

LUVs were prepared from the desired lipid mixture: POPC, POPC:Chol (67:33 molar ratio), or DOPC:Chol:SM (37:46:17 molar ratio). For immobilisation on the glass surface and lipid diffusion measurements, lipid derivatives DSPE-PEG-biotin and Atto647N-DPPE were included, each at approx. 1 molecule per 1000 lipids, whereas for GP experiments C-Laurdan was added at the same concentration. All lipid stock solutions were prepared in chloroform and stored at −20 °C. LUVs of heterogeneous sizes (unextruded LUVs) were formed by drying the lipid mixture by evaporating the organic solvent in a vacuum desiccator for 1 h and its rehydration in PBS while vortexing vigorously for 10 min, yielding a suspension of approx. 0.5 mM. To obtain LUVs of uniform sizes approx. 200 nm in diameter, part of the heterogeneous vesicle suspension was passed through 200-nm pores (Whatman) 20 times using a manual mini-extruder (Avanti Polar Lipids) preheated to 45 °C. Vesicles were stored at 4 °C and used in experiments within two days.

### Preparations of Supported Lipid Bilayers (SLBs)

SLBs were prepared by spin-coating, as previously described [13], using the same lipid mixtures as for LUV preparations, but without the biotinylated lipid and at a lower concentration of the fluorescent lipid probe (1 molecule per 10^4^ lipids). Every solution of lipids in chloroform and methanol (1:1 volume ratio, 1 mg lipids/ml) was spin-coated on a piranha solution-cleaned round 25-mm coverslip (thickness #1.5, VWR) for 45 s at 3200 rpm and rehydrated with SLB buffer (10 mM HEPES, 150 mM NaCl, pH 7.4). Prepared SLBs were kept hydrated in the SLB buffer and used immediately for the measurements.

### LUV immobilisation for STED-FCS measurements

For microscopy and FCS experiments, LUVs containing the biotinylated lipid were immobilised in 8-well glass-bottomed Ibidi chambers exploiting the biotin-streptavidin-biotin sandwich linker. This approach prevents vesicle flattening and rupture and does not influence their lipid membrane mobility [14]. For this purpose, the chambers were coated with a mixture of BSA and biotinylated BSA (5:1 molar ratio, 1 mg/ml) for 1 h, washed several times with PBS, incubated with streptavidin (500 ng/ml) for 1 h, and washed with PBS again multiple times. Thereafter, ten-fold diluted PBS suspension of LUVs was added in the prepared chambers for incubation for approximately 30 min. Finally, the non-adhered LUVs were removed by carefully washing the chambers with PBS prior to measurements.

### Acquisition of TCSPC data

Experiments were performed at room temperature using a Leica SP8 STED instrument equipped with a 100x/1.4 NA oil immersion STED objective. The lipid probe Atto647N-DPPE was excited by the 633-nm line from the white light laser pulsing at 80 MHz (average power 0.2 or 0.6 μW), depleted with a doughnut-shaped 775-nm pulsed STED laser (average power 55 mW), and recorded with a hybrid detector in the wavelength range 640–730 nm. Under these conditions, a 3-fold improvement in lateral resolution with respect to confocal was achieved (estimated from the ratio of transit times of freely diffusing fluorescent lipids in POPC SLBs in confocal vs. STED mode, being 3.6 and 0.4 ms, respectively) [13], resulting in an effective observation spot of approx. 75-nm in diameter. Within each sample, confocal and STED imaging was used to select vesicles of comparable brightness and size: around 0.2 and 0.5–1 μm in diameter for the extruded and non-extruded LUVs, respectively. Time-correlated single photon-counting (TCSPC) streams from the sites of selected LUV were acquired for 10–30 s using PicoQuant Hydraharp 400 electronics and SymPhoTime software.

### Analysis of STED-FCS data

The recorded TCSPC streams were gated to minimise the effects of scattered light using FoCuS-point software [15], and intensity time traces at frequency 100 kHz were generated. These were then cropped to the regions with the signal above the background level (typically yielding traces of 2–20 s, depending on the excitation power as well as lipid composition and size of LUV), and corrected for photobleaching using the local-averaging method over 0.25–0.5 s intervals, as implemented in the FoCuS-scan software [16]. FoCuS-scan software was used also to calculate the correlation functions and fit them with a 2D diffusion model:

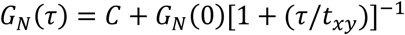

where *G_N_(τ)* is the correlation function at time lag *τ*, *C* the offset, *G_N_(0)* the amplitude, and *t_xy_* the average lateral transit time through the observation spot. From the obtained lateral transit times, diffusion coefficients were calculated using the known diameter of the STED-FCS observation spot (75 nm, see above). Time traces with artefacts due to vesicle movement, and FCS curves with low signal-to-noise ratio or high fit parameter errors, were discarded based on single-datapoint evaluation, prior to any comparison to avoid bias.

### Generalised Polarisation (GP) measurements

Emission spectra of LUV suspensions were measured in transparent-bottom 96-well plates (Porvair) by a microplate reader (Clariostar), which excited the samples at 385 nm. To calculate the generalised polarization values (GP) of the background-subtracted spectra, fluorescence emission intensities at 440 nm (I_440nm_) and 510 nm (I_510nm_) were used:

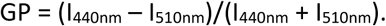

## 3. Results

To determine the effect of the membrane curvature and lipid composition on molecular mobility on the virus surface, we have utilised a synthetic lipid vesicle system. LUVs were generated using either POPC only (LUV-Ld), POPC:Chol mixture at 67:33 molar ratio (LUV-Lo), or DOPC:Chol:SM at 37:46:17 molar ratio (LUV-HIV). POPC vesicles represent the single component highly disordered and fluid lipid membrane, while POPC:Chol mixture represents a more rigid and ordered lipid composition. The POPC:Chol:SM mixture represents a simplified synthetic HIV-like lipid mixture with the similar GP value and molar ratio of Chol and SM to those found in the real virus [7]. For measurements of lipid mobility in membranes with different curvatures, LUVs of two size classes were selected – diameter around 0.5–1 μm, or around 0.2 μm. The latter were generated by extrusion through 200 nm pore to obtain vesicles with similar size to the HIV-1 virus (Figure 1 a).

To analyse the lipid mobility in each of these conditions, LUVs were generated through the addition of a fluorescent lipid analogue DPPE-Atto647N (1:1000 ratio) and were immobilised on glass coverslips (Figure 1 b) using biotin-streptavidin-biotin sandwich linker [14]. In this study we use fluorescence correlation spectroscopy (FCS) to investigate lipid diffusion. In FCS, diffusion coefficients of fluorescently labelled molecules are quantified by analysing fluctuations in the fluorescence signal as those molecules diffuse in and out of the microscope’s observation spot [17,18]. To realize such measurements on vesicular structures smaller than 250 nm (i.e. smaller than the observation spot of the conventional confocal microscope), we are employing FCS on a super-resolution STED microscope capable of observation spots < 100 nm (STED-FCS) [11,12]. In our current STED-FCS measurements, the STED microscope was tuned to yield an effective diameter of the observation spot of around 75 nm (see Materials and Methods), thus well below the size of the smallest measured vesicle. Due to the low copy number of Env proteins per individual virus, our previous study of Env mobility on HIV-1 surface utilised scanning STED-FCS (sSTED-FCS) [6], which minimizes photobleaching and enables accurate recovery of diffusion coefficients for slowly moving molecules only (such as Env or other proteins in the viral membrane). The number of fluorescent lipid analogues per individual LUV is much higher, thus making photobleaching a less critical issue. Consequently, we have used point STED-FCS in this case, offering the high temporal resolution needed to follow 1000-fold faster diffusion of lipids in LUV compared to protein diffusion in viral membranes. Photobleaching was still minimized by a very low excitation power (10–25-fold lower compared to the sSTED-FCS study [6]).

Lipid mobility data was acquired for individual 200 nm or ~1 μm fluorescent vesicles using point STED-FCS in time-correlated single photon-counting mode (TCSPM). TCSPM mode was used in order to provide additional fluorescence lifetime-based filtering that allows for removal of residual laser scattering thus increasing signal-to-noise ratio of the acquired signal [19,20]. In addition, photobleaching correction was also applied using a local-averaging method [16]. The resulting autocorrelation curves (Figure 1 c) were fitted with a generic two-dimensional (2D) diffusion model to obtain the average transit times of fluorescent lipids through the subdiffraction sized observation spot and to derive the diffusion coefficient for each sample.

Firstly, we compared LUV lipid mobility with the behaviour of the same lipid mixes in supported lipid bilayers (SLB, a lipid bilayer spin-coated onto the microscope cover glass) – a standard model for the measurements of lipid mobility (Figure 2). Results showed that in all LUV formulations the recorded lipid diffusion coefficient was consistently higher (~2-3 fold) than the corresponding SLBs with the same lipid composition. This consistent difference is not caused by the incomplete modelling of the molecular diffusion in spherical lipid surfaces by a simple 2D diffusion equation; we have in our previous study of HIV-1 surface mobility shown that more sophisticated fitting models that consider the diffusion on the surface of a spherical vesicle give comparable results [6]. Similar 2-fold diffusion coefficient differences between giant unilamellar vesicles and SLBs with similar lipid compositions were previously observed [21]. Most likely this reduction in lipid mobility observed for supported SLBs compared to free-standing vesicular membranes is caused by a slow-down of lipid diffusion due to van der Waals interactions of lipid headgroups with the glass surface [22]. Nevertheless, the consistency of our results between samples of different lipid composition indicates that the diffusion coefficients obtained for the LUVs represent a good estimate of the molecular mobility in this model system.

**Figure 2.**
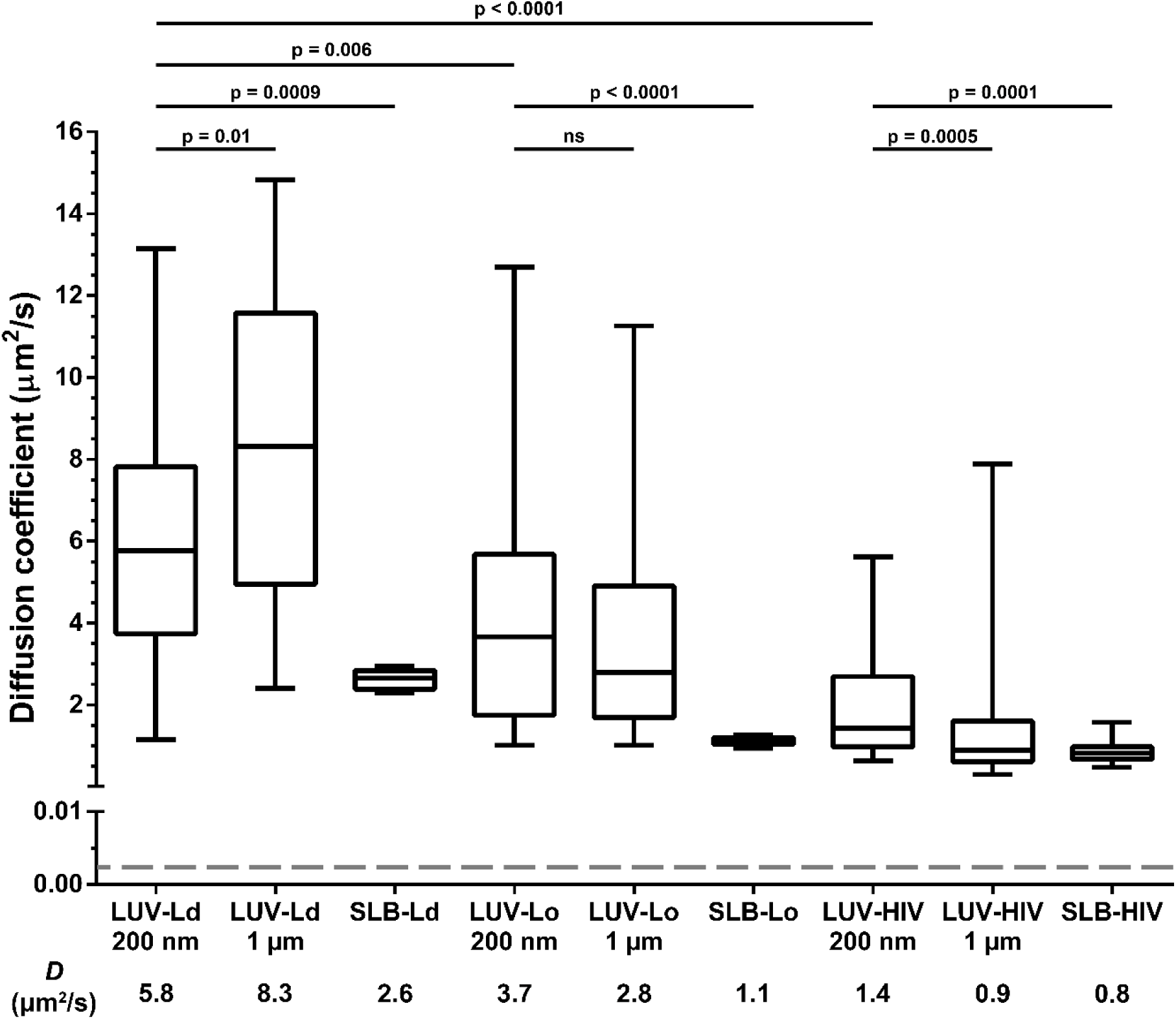
Diffusion coefficients of a fluorescent lipid DPPE-ATTO647N in 200 nm and 1 μm sized Large Unilamellar Vesicles (LUV) and Supported Lipid Bilayers (SLB) with the following compositions: POPC (LUV-Ld, SLB-Ld), POPC:Chol at 67:33 molar ratio (LUV-Lo, SLB-Lo) and DOPC:Chol:SM at 37:46:17 molar ratio (LUV-HIV, SLB-HIV). Median diffusion coefficients (D) were determined by STED-FCS measurements of 40 LUVs and 5 SLB positions each from two independent measurements. Results are shown in a box (IQR) and whisker (min/max) plot and the statistical significance was assessed by Wilcoxon rank-sum test. Grey dashed line represents surface Env diffusion coefficient determined for mature HIV-1 particles [6].

We then compared the impact of the lipid composition and vesicle diameter on the lipid diffusion coefficients in LUVs (Figure 2). Comparison of the diffusion coefficients for 200 nm and 1 μm vesicles for different LUV formulations shows varying relations: the reduction in LUV size caused a decrease in diffusion coefficient for POPC membranes (*D* = 5.8 compared to 8.3 μm^2^/s), no significant difference for the POPC:Chol LUVs (*D* = 3.7 compared to 2.8 μm^2^/s), and an increase for the POPC:Chol:SM mix (1.4 compared to 0.9 μm^2^/s). On the other hand, the comparison of the LUVs of the same size but different compositions of increasing rigidity and similarity to the HIV-1 lipid envelope (from LUV-Ld to LUV-HIV) shows consistent and significant reductions of the diffusion coefficients (*D* = 5.8 μm^2^/s for LUV-Ld, 3.7 μm^2^/s for LUV-Lo, and 1.4 μm^2^/s for LUV-HIV in the case of the 200-nm sized LUVs). These results identify the lipid composition of HIV-1 membranes, rather than their curvature, as one of the major factors responsible for the reduction of mobility of the molecules on HIV-1 surface. Considering that the membrane curvature of these particles is, in comparison to the membrane thickness of approx. 4 nm, still relatively mild (see insets in Figure 1 a), stronger effects of the curvature are expected, and indeed observed, at even smaller sizes below 50–100 nm [23–25].

To further confirm the importance of increased lipid packing, but not the curvature of the vesicle membranes in the modulation of lipid envelope mobility, we measured emission spectra of a polarity-sensitive dye C-Laurdan [26] in the membranes of all LUV compositions and sizes (Figure 3 a). The extracted GP values for POPC vesicles were similar to values reported previously [5]. Moreover, extruded LUVs consistently showed red-shifted spectra and thus lower GP values (~0.07 unit difference) than corresponding larger unextruded LUVs, indicating, on average, less dense lipid packing compared to larger vesicles. However, the differences in GP values were much larger for different lipid compositions than for different sizes within each composition. In fact, increasing GP values (indicative of denser lipid packing) appear well correlated with a decrease in LUV lipid diffusion coefficients (Figure 3 b).

**Figure 3.**
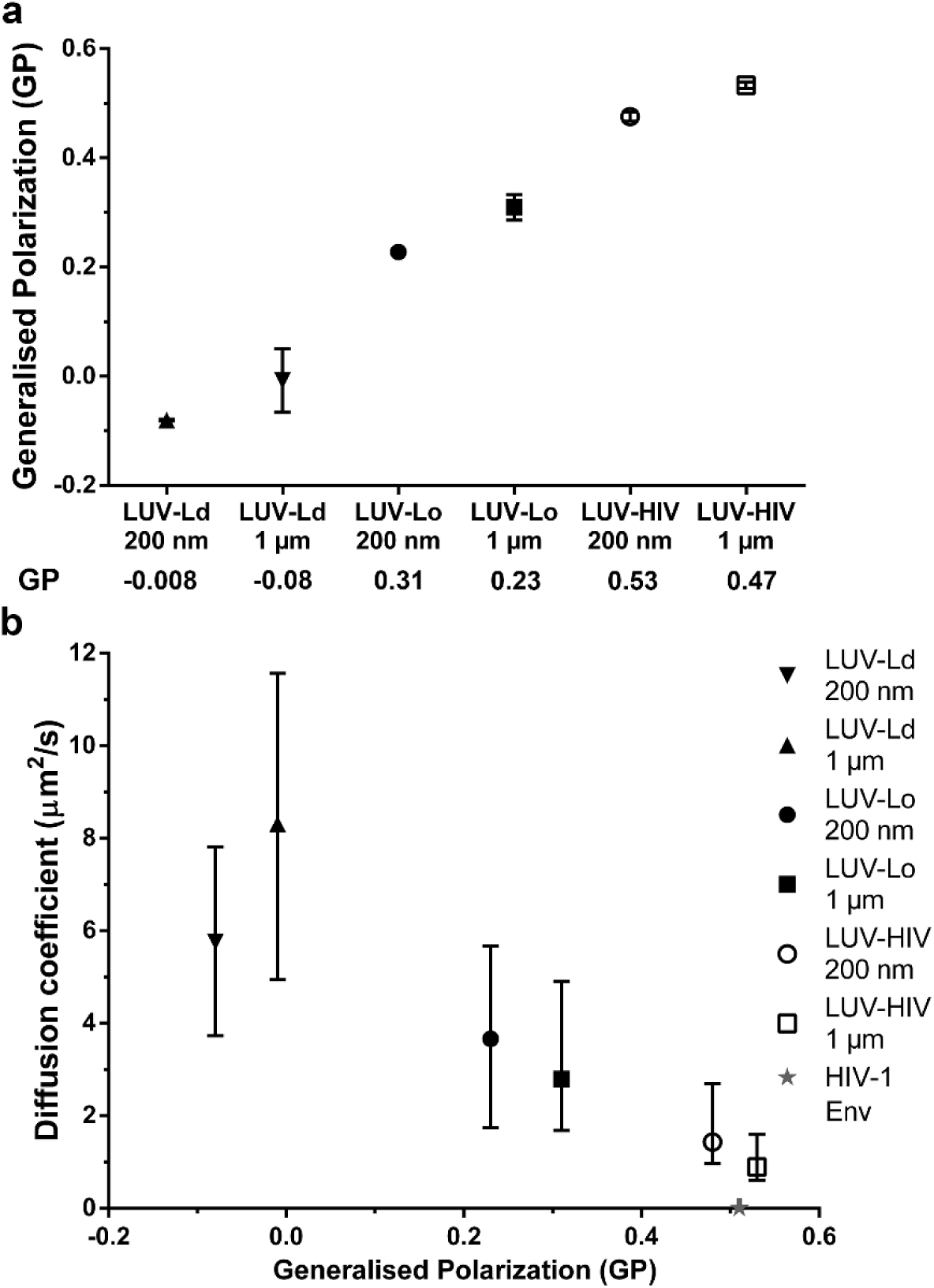
Analysis of LUV membrane properties by C-Laurdan staining. (**a**) Mean ± SD values of the generalised polarisation (GP) parameter were determined for 200 nm pore size extruded or 1 μm unextruded LUV suspensions with following compositions: POPC (LUV-Ld), POPC:Chol at 67:33 molar ratio (LUV-Lo) and DOPC:Chol:SM at 37:46:17 molar ratio (LUV-HIV). Results represent data from two independent measurements; (**b)** Median diffusion coefficient vs average GP values plotted for all LUV sizes and compositions. Error bars represent IQR of the LUV diffusion coefficients. Grey star represents Env diffusion coefficient and GP values in mature HIV-1 particles [6].

## 4. Discussion

Little is known about the dynamic properties of the HIV-1 virus surface. This study refines the findings of the previous work that described the low mobility nature of HIV-1 lipid envelope [6]. By measuring diffusion coefficients of lipids and membrane polarity of LUVs of different sizes and compositions, we herein established that the tightly packed nature of HIV-1 lipid envelope, rather than its high curvature, plays a major role in the creation of a very low mobility environment on the virus particle surface.

However, despite observing 4-fold reduction in lipid mobility between most fluid and most rigid LUV compositions, it is clear that this factor is only part of the reason for the even further 700-fold lower molecular mobility of proteins previously observed in *bona fide* HIV-1 particles (as exemplified by a dashed line and a star in Figures 2 and 3b, respectively; unfortunately, we were unable to measure the diffusion of fluorescent lipid analogues in HIV-1 particles also due to their efficiency of incorporation being too low). Even accounting for the fact that proteins diffuse approximately 10-fold slower in cell membranes than do lipids [6,11], this difference alone is insufficient to explain the extremely low mobility of Env observed in HIV-1 membrane (*D* ≈ 0.002 μm^2^/s). This discrepancy may be explained by other features of HIV-1 particles that are not present in LUVs such as tightly packed internal virus structures [27], lipid-MA protein interactions [28] and incorporation of Env as well as a large variety of cellular proteins on the virus surface during the budding process [29].

Our study has provided additional insights into the dynamic properties of HIV-1 surface. It further highlighted the importance of rigid and ordered lipid composition as the determinant of the dynamic behaviour of molecules on HIV-1 virus surface [6], which, in turn, was previously shown to underscore the ability for HIV-1 to successfully fuse with the target cell [10]. Future studies of this still relatively unexplored aspect of HIV-1 replication cycle may provide a novel therapeutic approach that could potentially prevent virus entry by subtle alterations in lipid packing of the HIV-1 envelope.

## Author contributions

JC, IU and CE conceived the study and designed the experiments. IU, JB and DS performed the experiments. IU analysed the data. DW provided technical support for the analysis of the data. IU, JC and CE wrote the paper.

## Acknowledgements

We would like to thank the Wolfson Imaging Centre (Weatherall Institute of Molecular Medicine, Oxford), especially Christoffer Lagerholm for help on the microscopes. We acknowledge support by the MRC (grant number MC_UU_12010/unit programs G0902418 and MC_UU_12025), MRC/BBSRC/EPSRC (grant MR/K01577X/1), Wellcome Trust (grant 104924/14/Z/14), Deutsche Forschungsgemeinschaft (Research unit 1905 “Structure and function of the peroxisomal translocon”), and Oxford internal funds (EPA Cephalosporin Fund and John Fell Fund). IU is funded by a Marie-Skłodowska-Curie individual fellowship (grant 707348). JB is supported by Wellcome Trust PhD Programme (203853/Z/16/Z).

## Conflicts of interest

The authors declare no conflict of interest.

